# Functional and phylogenetic β diversities and their link with clustering/overdispersion and uniqueness/redundancy

**DOI:** 10.1101/2023.09.01.555888

**Authors:** Sandrine Pavoine, Carlo Ricotta

**Affiliations:** Centre d’Ecologie et des Sciences de la Conservation (CESCO), Muséum National d’Histoire Naturelle, CNRS, Sorbonne Université, 75005, Paris, France; Department of Environmental Biology, University of Rome ‘La Sapienza’, Rome, Italy

**Keywords:** biodiversity measure, biodiversity conservation, community dissimilarities, diversity partitioning, ecological and evolutionary processes, ecosystem functions, Hill’s numbers, processes, quadratic entropy, resilience

## Abstract

In the recent decades, research on biodiversity in community ecology has been marked by the consideration of species’ evolutionary histories and biological traits, including those said functional as they relate to the functions a species has in ecosystems. Among the different spatial levels at which FP diversity can be quantified, less attention has been given to the definition of the general concept of between-communities (β) FP diversity than to those of local, within-communities (α) and regional, merged-communities (γ) FP diversities. Here we develop a new way for partitioning FP β diversity into elementary components to underline how and why FP β diversity differs from species β diversity, the latter reflecting only differences in species’ abundances between communities. As a reference example, we consider two distinct measures of FP β diversity: Rao’s dissimilarity coefficient (*Q*_*β*_) that expresses an average dissimilarity between communities, and its transformation into an equivalent number of communities (*D*_*β*_).

Thanks to analytical partitioning and simulations, we show that *Q*_*β*_ and *D*_*β*_ are connected differently with typical patterns of community structure. The search for the ecological and evolutionary processes that drive community assembly and the assessment of community resilience and stability have indeed revealed typical community structures: the local clustering of species with similar traits or shared evolutionary histories; and the local (α) or regional (γ) presence of functionally or phylogenetically redundant versus unique species. While *Q*_*β*_ and *D*_*β*_ are both increasing functions of species β diversity and γ FP uniqueness, *Q*_*β*_ increases with FP clustering while *D*_*β*_ increases with α FP redundancy. The selection of an index of β diversity for a given study depends on its objective, such as communication to conservation policy makers or theoretical study of the ecological and evolutionary processes of community assembly. To facilitate and secure this selection, we call, through our study, for the development of formal and precise definition(s) for FP β diversity in light of the concepts of clustering versus overdispersion, and redundancy versus uniqueness. In particular, we call for further research on when and why FP β diversity should increase with FP clustering.

## INTRODUCTION

In the context of the current biodiversity crisis, the need has never been so crucial to understand how the current amount of biodiversity in different parts of our planet varies (e.g., Albert et al., 2021; Borges et al., 2020; Davis et al., 2018). A prerequisite step for such an understanding is the development of appropriate ways to measure biodiversity (Table 1). The interspecies component of biodiversity focuses on the number of species and their relative abundances and on the variety of their biological characteristics such as their traits (=well-defined, measurable phenotypic properties) and phylogeny (=relationships between species reflecting evolutionary history). In community ecology, the key unit where biodiversity is quantified is a community: set of interacting populations of the species living within an area at a particular time. Since Whittaker (1972), biodiversity has often been studied within communities (=α diversity), between communities (=β diversity) and over all communities (=γ diversity). This allowed researchers to analyze ecological and historical processes that drive biodiversity patterns while acting either at a local level or at a regional level (e.g., Pavoine & Bonsall, 2011). In real case studies, both α and β diversities contribute to the global γ diversity of the whole set of communities. Despite all scales of biodiversity (α, β, γ) are critical to maintain multiple ecosystem functions and services (Pasari et al., 2013), the β component has been the least studied among the myriad of studies on biodiversity, and is the least precisely defined. The variation in species composition among communities it represents is yet an important insurance for the maintenance of ecosystem multifunctionalities and services at a landscape level (Mori et al., 2016). The challenge for ecologists has thus been to quantify each component of diversity (α, β, γ) and the measurement of the β component raised some debates (e.g., Jost, 2006; Jurasinski et al., 2009; Schmera et al., 2020; Veech & Crist, 2010).

**TABLE 1.** Glossary.

| Concept | Definition |
| --- | --- |
| $\alpha$ diversity | Amount of differences between individuals within communities |
| $\beta$ diversity | Amount of differences between communities |
| $\gamma$ diversity | Amount of differences between individuals both within and between communities |
| FP* clustering | Clustered communities are different from each other but individuals within communities are similar. |
| FP* overdispersion | Overdispersed communities are similar from each other and individuals within a community have different biological characteristics. |
| FP* randomness | In case of randomness, individuals are distributed among the communities randomly as regards their traits or phylogenetic positions. |
| $\beta$ redundancy | $\beta$ redundancy occurs when communities have different species but for each species in a community one can find a similar one with similar biological characteristics in each other community. |
| $\beta$ uniqueness | $\beta$ uniqueness occurs when trait-based or phylogenetic $\beta$ diversity mimics species $\beta$ diversity. |
| $\alpha$ and $\gamma$ redundancy | Redundancy at the $\alpha$ (resp. $\gamma$) scale means that individuals within a community (resp. all individuals over all communities) are similar. |
| $\alpha$ and $\gamma$ uniqueness | In contrast, uniqueness at the $\alpha$ (resp. $\gamma$) scale means that species are highly different from each other within a community (resp. all individuals over all communities). |

\* FP = (trait-based,) functional or phylogenetic.

Explaining why the debates were raised is easier than solving them. The debates stem from the fact that biodiversity is a complex, multidimensional concept that we try to quantify by a single value or a few values. They also stem from the fact that biodiversity is quantified, even in ecology, for numerous objectives. Three main nonexclusive objectives can be cited here: “Obj1” saving biodiversity for its intrinsic value, “Obj2” maintaining ecosystem functioning and services, and “Obj3” understanding the processes underlying biodiversity patterns. When Obj1 is of interest, the main challenge for developing measures of biodiversity is to count all elements of as many aspects of biodiversity as possible without taking the risk of missing and losing a facet of biodiversity. When Obj2 is of interest, the main challenge is to identify which aspects of biodiversity enable stable ecosystem functions. When Obj3 is of interest, the main challenge is to disentangle the different components of biodiversity to succeed in explaining how and why each component varies in space and time. Clearly the different objectives call for different ways of quantifying biodiversity.

More generally, while α and γ diversities are generally well defined, β is often relegated to a function of γ and α: knowing that α and β both contribute to γ, it has been implicitly admitted that, in γ diversity, what is not α is β yielding γ=α+β (additive partitioning) or that β is the ratio of γ to α, so that γ=α×β (multiplicative partitioning). In the former case, α and β are considered as a partition of γ, and they are measured with the same unit; while in the second case they have distinct units. The exact meaning of the component of β diversity is however often ambiguous (Ricotta et al., 2010) so that frameworks that justify and organize β diversity measures still have to be developed (e.g., Tuomisto, 2010a,b). Jurasinski et al. (2009) referred to such definitions of β as “proportional diversity” because their associated measure results from the comparison of diversity measurement at two embedded spatial levels (α and γ) rather that being the direct resultant of measures of differentiations in the compositions of the communities.

Although α, β and γ are intrinsically linked as exemplified by partitioning approaches of γ into α and β, an apparent consensus seems to have been reached on the fact that the range of variation of β must be independent of α diversity (Chao et al., 2012). Within this range of variation, β diversity may, however, be a function of α diversity among other sources of variation. We name this property the range-independence from α diversity (RIαD). Indices that do not fulfill this property were thus rejected and transformed into other indices once named “true diversity” or Hill numbers (Jost, 2006; but see Hoffman & Hoffman, 2008) that fulfills the RIαD property (e.g., Chiu et al., 2014; Ricotta & Szeidl, 2009). With “Hill numbers” indices, γ is thus the resultant of two components α and β with independent ranges of variation, while the components themselves are not statistically independent as each individual in each community contributes to both α and β diversities (see also, Veech & Crist, 2010). This partial independence and the fact that even indices satisfying the RIαD property can be “proportional diversity” indices (Jurasinski et al., 2009) imply that the question of the meaning of β diversity also concerns “Hill numbers” indices.

Roughly speaking, species β diversity represents the differences in species presences and/or abundances between several communities. When more information on species are available such as their relative phylogenetic positions or their traits, including traits said functional (as they determine species’ response to environmental factors or species’ effect on ecosystem functioning, Lavorel & Garnier, 2002), then β diversity can reflect differences in the phylogenetic or trait-based compositions of the communities. While trait-based β diversity may reveal local ecological processes, phylogenetic β diversity could give insights on global dispersal, speciation and extinction events. Hereafter we will refer to these different aspects as FP for “functional or phylogenetic”. With the addition of these FP aspects of diversity in community ecology, many research studies in the last two decades have also tackled the concepts of clustering and overdispersion (Webb et al., 2002; Weiher & Keddy, 1995). If species within communities are similar, having similar traits or phylogenetic positions while species in distinct communities are different, having different traits or distant phylogenetic positions, then species are said clustered within communities. This pattern is referred to, here, as the FP clustering. If, on the contrary, species within communities are different while for each species in a community there exists another species in another community with similar traits or a close phylogenetic position, then species are said overdispersed within communities. This pattern is referred to, here, as the FP overdispersion. When individuals disperse and colonize communities independently of their traits or phylogenetic position, then communities are randomly assembled, regarding their FP characteristics.

The interest in clustering and overdispersion patterns was raised, in the context of Obj3, by the willing of finding the ecological and evolutionary processes that drive community assembly. Notably, trait-based clustering was associated with environmental filtering (if traits are related to the environmental niche) and exclusion of poor competitors (if traits are related to fitness, Mayfield & Levine, 2010). Trait-based overdispersion has been mostly associated with limiting similarity that would avoid niche overlaps and competition (e.g., Bañares-de-Dios et al., 2020; Cavender-Bares et al., 2004; Ramm et al., 2018; but see Gallien, 2017) but can also result from other processes. For example in plant communities it can result from colonization of distant relatives in late successional stages (Li et al., 2015). Clustering and overdispersion are often evaluated by comparing the observed value of FP α diversity with values expected under a null model of species distributions with a constrained, fixed γ diversity (e.g., Cooper et al., 2008; González et al., 2022; Lanier et al., 2013; Webb et al., 2002). When β is a simple function of γ and α such as γ-α or γ/α and knowing that γ is fixed under the null model, the same test can equivalently be defined using FP β rather than α diversity: observed β > average simulated β (under null model) expresses a trend of clustering over the whole studied area, while observed β < average simulated β expresses a trend of overdispersion. How the component of β diversity directly relates to these concepts of clustering and overdispersion has, however, not been formally explained so far.

In the context of Obj2 and Obj3, comparing FP diversity patterns to species diversity patterns at the α and γ levels may both help in identifying assembly rules (Pavoine & Bonsall, 2011) and in assessing the vulnerability of ecosystem functioning and services (Biggs et al., 2020). One way to make this comparison is to measure uniqueness (U) indices as the ratio of FP diversity to species diversity and redundancy (R) indices as inverse of uniqueness indices (R = 1/U). Note that the maximum of FP diversity can be made equal to species diversity if one considers that species diversity represents a special case where species are maximally dissimilar in their traits or phylogenetic positions (Ricotta et al., 2020). This requires a restriction on the use of diversity indices: that species diversity is higher than FP diversity. This property was named the “redundancy property” (Appendix S1 in Ricotta et al., 2020). Well-used α and γ diversity indices respect this property. As a result, a range of α and γ redundancy indices can easily be developed also as complement of uniqueness indices (R = 1-U). High FP diversity combined with high FP redundancy could ensure the resilience and stability of ecosystem functioning and associated services for human well-being if high FP diversity ensures ecosystem multifunctionalities and high species diversity compared to FP diversity (i.e. high FP redundancy) permits that the effect of any species loss by disturbance can be mitigated by the presence of other species that can ensure similar functions. Recently Ricotta et al. (2020) developed the complementary concept of β uniqueness as the ratio of FP β diversity to species β diversity. However how β diversity directly relates to the concepts of α and γ uniqueness and redundancy is still an open question.

The research on “independent” α and β components of diversity emerged from the observation of low values of β diversity, notably when the Gini-Simpson index of species diversity (Jost, 2006) or its generalization, Rao’s quadratic entropy, for FP diversity (Ricotta & Szeidl, 2009) were used, even when differences could be observed between the communities. Rao’s quadratic entropy has also been frequently used in association with null models to detect FP clustering and overdispersion (e.g., Ren et al., 2023). The development of β redundancy indices that compare β FP diversity with β species diversity also started with Rao’s quadratic entropy (Ricotta et al., 2020). Rao’s quadratic entropy is the average abundance-weighted FP dissimilarity between species in a community. It is well used (e.g., Chen et al., 2019) and a reference to measure diversity in community ecology. We thus explored Rao’s quadratic entropy and one of its transformation into Hill numbers, to evaluate the meaning of its component of β diversity, in link with species β diversity, redundancy and uniqueness, clustering and overdispersion.

## METHODS

### α, β, γ diversities

#### The initial apportionment by C.R. Rao

Consider *M* areas that represent *M* species communities, numbered 1 to *M*. Let *S*_*m*_ be the number of species present in area *m*, for a total of *S*_*γ*_ different species over the whole set of areas 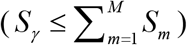 be the matrix of FP dissimilarities between species (*d*_*ii*_ = 0, *d*_*ij*_ = *d*_*ji*_). Let 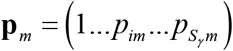 be the vector of relative abundances of the *S*_*γ*_ species in area *m* (species absent from area *m* have an abundance equal to zero). Let *w*_*m*_ be a weight attributed to area 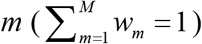 the global relative abundance of any species *i* over all areas is 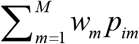.

The value of the quadratic entropy applied to area *m* is

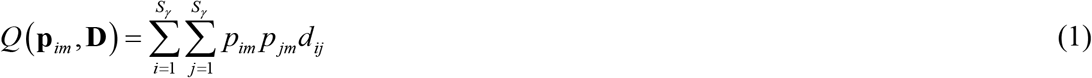

Rao (1982) defined the diversity within areas (α diversity) as (see detailed equations in Appendix S1) 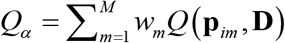, the global diversity over all areas (γ diversity) as 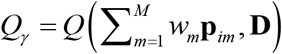 and the dissimilarity coefficient (DISC) between areas (β diversity) as *Q*_*β*_ = *Q*_*γ*_ − *Q* . The for a of *Q*_*β*_ diversity means that in γ diversity what is not α diversity is β diversity. It can also be viewed as the quadratic entropy applied to the weights (*w*_*m*_’s) attributed to areas and to FP dissimilarities between areas. Let **w** be the vector of *w*_*m*_ values and Δ=(*δ*_*mn*_) be the matrix of FP dissimilarities between areas with

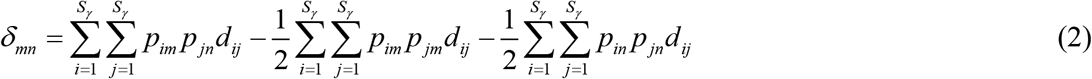

Then *Q*_*β*_ = *Q* **(w**, Δ) . The dissimilarity between any two areas *m* and *n* is thus measured in *Q*_*β*_ by the difference between the average FP dissimilarity between one species drawn from area *m* and the other drawn from area *n* and the average FP dissimilarity between two species drawn from the same area (either *m* or *n*). This formula satisfies the condition that the dissimilarity between two identical areas is null. In contrast an alternative formula that would only use, as a measure of FP dissimilarity between areas *m* and *n*, the average FP dissimilarity between one species drawn from area *m* and the other drawn from area *n* (as, e.g., in Webb et al., 2008) would not satisfy this elementary condition (Ricotta et al., 2015).

The equation *Q*_*β*_ = *Q* **(w**, Δ) exemplifies that, in Rao’s apportionment of quadratic entropy, species and communities are treated in the same manner. The whole γ diversity depends on the relative abundance of species and the FP dissimilarities between them and on the weights defined for communities and the FP dissimilarities defined between them.

If the *d*_*ij*_ are multiplied by a constant say λ, then *Q*_*β*_ is also multiplied by this constant. This means that *Q*_*β*_ depends on the global level of biological differences between species. For *Q*_*β*_ to have nonnegative values, the *d*_*ij*_s need to satisfy mathematical properties named the Euclidean properties (= the distance values 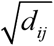 form a cloud of points in a Euclidean space where each point is a species). Without any additional restrictions on the definition of the *d*_*ij*_ values, the range of possible values for *Q*_*α*_, *Q*_*β*_, and *Q*_*γ*_ is [0, ∞[. The range of *Q*_*β*_ is independent of *Q*_*α*_ (i.e., *Q*_*β*_ satisfies the RIαD property).

#### The concept of maximally dissimilar species

Applications of the quadratic entropy, however, generally revealed an additional restriction on the choice of the *d*_*ij*_: 0 ≤ *d*_*ij*_ ≤ 1, meaning that FP dissimilarity between species are bounded (Ricotta & Szeidl, 2009). We thus hereafter consider this restriction: the maximum value of *d*_*ij*_ is unity. The case *d*_*ij*_ = 1 means that species *i* and *j* are “maximally dissimilar”. Consider a set of *S* species, with this restriction, the maximum possible value of the quadratic entropy applied to this set, over all matrices of distances between species and all vectors of species abundance, is max**p**,**D {***Q* **(p, D)}** = (*S* −1) / *S* . It is reached when species have equal abundances (*p*_*i*_ = 1/S for all species *i*) and are all maximally dissimilar from each other (*d*_*ij*_ = 1 for all *i*≠*j*). This maximum tends to 1 when *S* increases. When the FP dissimilarities are bounded so that there exist maximally dissimilar species, the value of the quadratic entropy is thus bounded between 0 and 1, and this applies both to the α and γ levels of diversity. As a consequence, for a given value of *Q*_*α*_, *Q*_*β*_ is bounded between 0 and 1 - *Q*_*α*_: hence, in that case, the range of *Q*_*β*_ depends on *Q*_*α*_.

Ricotta and Szeidl (2009) proposed, in this particular case of bounded FP dissimilarities, to use a modification of the quadratic entropy and a new apportionment scheme so that a component of β diversity can be defined with a range independent from α diversity. Let *s*_*ij*_=1-*d*_*ij*_ be the FP similarity between species. It yields the following component of α diversity

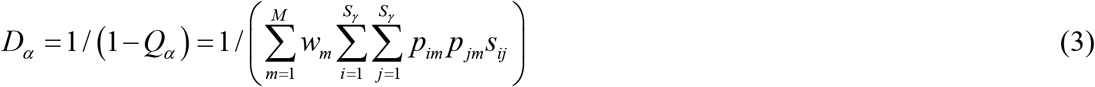

the following global diversity over all areas (γ diversity)

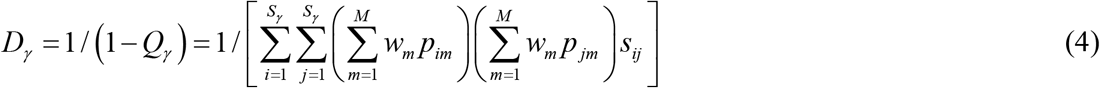

and the following dissimilarity coefficient between areas (β diversity)

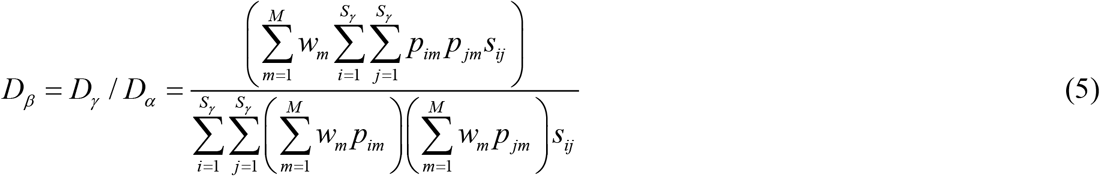

which is the ratio of the average FP similarity at the α level to the average FP similarity at the γ level. While *Q*_α_ and *Q*_γ_ represent the average dissimilarity between species at the α and γ scales, respectively, *D*_α_ and *D*_γ_ represent the inverse of the average similarity between species at the α and γ scales.

*Q*_*β*_ represents the gap between the dissimilarity among individuals at the γ level and that at the α level. High values of *Q*_*β*_ are expected for high γ diversity and low α diversity. In contrast, *D*_*β*_ is an equivalent number of areas and varies thus between 1 and *M*. Its maximal value *M* is reached as soon as communities are evenly weighted (*w*_*m*_=1/*M* for all *m*) and maximally dissimilar (communities do not share species and *s*_*ij*_ = 0 if species *i* and *j* belong to distinct communities; whatever the global amount of diversity at the γ scale).

Note that transforming *Q*_*β*_ into 1/ (1− *Q*_*β*_) would also lead to an index of β diversity that varies between 1 and *M*. But this alternative transformation would not lead to an index with a range of variation independent from α diversity. In contrast, the *D*_*β*_ and *Q*_*β*_ are linked by the following equation

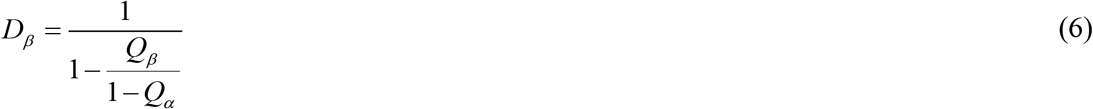

The standardization of *Q*_*β*_ by 1 − *Q*_*α*_in Eq. 6 generates the desired RIαD property that the range of variation of an index of β diversity is independent of α diversity.

#### Species-based β diversity

It is generally admitted that species-based indices consider that species are all maximally dissimilar from each other (*d*_*ij*_ = 1 for all *i*≠*j*). Let *Q*_*α*,spe.div_ *Q*_β,spe.div_ and *Q*_γ,spe.div_, *D*_*α*,spe.div_ *D*_β,spe.div_ and *D*_γ,spe.div_, be the values of *Q*_*α*_, *Q*_β_, *Q*_γ_, *D*_*α*_, *D*_β_, *D*_γ_, respectively, obtained by imposing that species are maximally dissimilar although they can vary in abundance. In that case (i.e. considering *d*_*ij*_ = 1 for all *i*≠*j*), *Q*_*β*_ and *D*_*β*_ reduce to

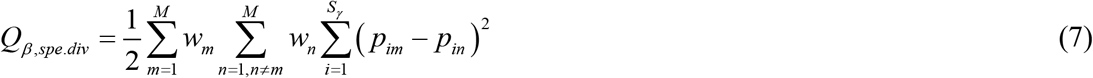

and

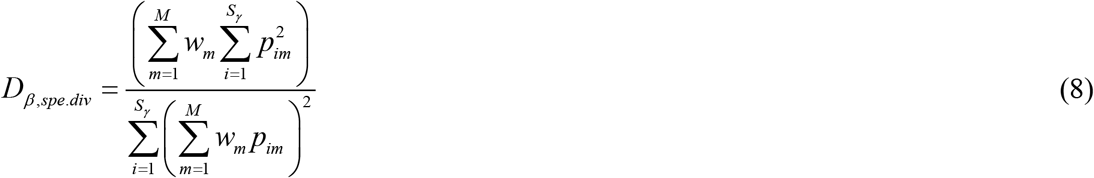

respectively. *Q*_*β*,*spe*.*div*_ is the average dissimilarity between any two areas, with this dissimilarity measured as the squared difference in species’ relative abundance between the two compared areas. It is the highest when in each area a species dominates in abundance but the dominant species in one area is different from the dominant species in the other area. *D*_*β*,*spe*.*div*_ is the ratio of the level of dominance within areas (α level) to the level of dominance over all combined areas (γ level). Here dominance is the degree to which a single species dominates in abundance, while others are rare. *D*_*β*,*spe*.*div*_ is the highest as soon as areas both do not share species and are evenly weighted (i.e., *w*_*m*_ = 1/*M*). When species are maximally dissimilar, *Q*_*β*_ and *D*_*β*_ are thus both high in case of clustering of identical individuals (=from the same species) within an area, while individuals are maximally different (=from different species) from one area to another. However, *D*_*β*_ is still high if individuals within an area belong to a range of distinct species, provided that individuals are maximally different from one area to another, while *Q*_*β*_ is not.

If in addition, species within areas have even abundances, we note *Q*_β,spe.ric_ and *D*_β,spe.ric_ the values of *Q*_β_ and *D*_β_, respectively, obtained by imposing that species are maximally dissimilar and have similar abundances within an area. We determined (proof in Appendix S1) that *Q*_*β,spe*.*ric*_ can be viewed as the proximity to a case of complete species clustering where there are as many areas as species, each area contains a single species and areas do not share species. *Q*_*β,spe*.*ric*_ is thus the highest in case of complete species clustering, while *Q*_*β,spe*.*div*_ is high both in case of complete species clustering (1 species per area), and when the species clustering is driven by uneven abundance distributions (in each area a species dominates in abundance but the dominant species in one area is different from the dominant species in the other areas). In contrast *D*_β,spe.ric_ is maximum as soon as areas do not share species (whatever the number of species in each area). *D*_β,spe.ric_ is related to a traditional family of dissimilarity indices that includes the Jaccard and Sørensen indices (e.g., Gower & Legendre, 1986; proof in Appendix S1).

This difference in behavior still exists when species are not maximally dissimilar. Indeed *Q*_*β*_ is high in case of clustering of similar individuals (=from species with close FP characteristics) within an area, while individuals are strongly different (=from different species with distinct FP characteristics) from one area to another. In contrast, *D*_*β*_ is high as soon as individuals are strongly different from one area to another as shown below.

#### Partitioning into elementary components

Below, we introduce partitioning schemes for *Q*_*β*_ and *D*_*β*_, where each of these indices of FP β diversity is compared to the value it would take if species were maximally dissimilar (i.e., if it simply reduced to species β diversity). Although the ratio of FP β diversity to species β diversity has been recently related to a concept of β uniqueness (Ricotta et al., 2020), this relatedness requires indices that respect the redundancy property: i.e., that species β diversity is higher than FP β diversity. Most β diversity indices do not respect this property, including *Q*_*β*_ and *D*_*β*_.

As shown in the previous section, the concept of clustering is crucial in the interpretation of *Q*_*β*_ values, while it is not in *D*_*β*_ values. To ease the interpretation of *Q*_*β*_ it would thus be useful to more explicitly depict how and how much clustering in FP characteristics determines levels of β diversity according to *Q*_*β*_. As values of *Q*_*β*_ are driven by FP clustering, one could simply envisage the ratio *Q*_*β*_ / *Q*_*γ*_ to measure the absolute level of FP clustering in a study area. Indeed, although the ratio of β to γ diversity has sometimes been used in the past as a measure of community dissimilarity, that is as a standardized measure to compare the compositions of communities (e.g., Hardy & Senterre, 2007; Lande, 1996), the ratio *Q*_*β*_ / *Q*_*γ*_ has also been interpreted as “local similarity excess”, that is as clustering (Hardy & Jost, 2008). High values of *Q*_*β*_ / *Q*_*γ*_ thus indicate high absolute level of FP clustering or equivalently high local FP similarity excess. However, high absolute FP clustering occurs as soon as individuals with similar functional trait values (from the same species) are clustered within an area, and sole members of the area. Maximum absolute FP clustering can thus be confounded with species clustering that occurs when individuals from the same species are clustered within an area, and sole members of the area or dominant in abundance in the area (species clustering = “local species identity excess” sensu Hardy & Jost, 2008).

To measure a relative level of FP clustering given the level of species clustering, one has to compare FP clustering with species clustering, yielding the following standardized coefficient:

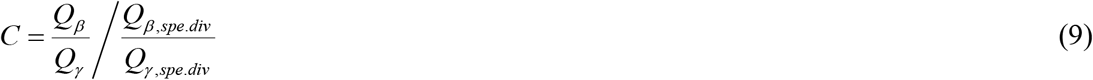

*C* higher than 1 means clustering due to FP characteristics of the species and *C* below 1 indicates overdispersion due to these characteristics. A typical example of case for *C* = 1 is obtained when the *d*_*ij*_ is constant over all species pairs *i,j. C* does not measure absolute clustering but relative clustering: the amount of FP clustering that can be attributed solely to the distribution of FP characteristics across areas.

*Q*_*β*_ can be apportioned into components of species-based β diversity (*Q*_*β,spe*.*div*_), global FP uniqueness (*U*_*γ*_ = *Q*_γ_/*Q*_γ,*spe*.*div*_) and FP clustering (*C*) using the following formula:

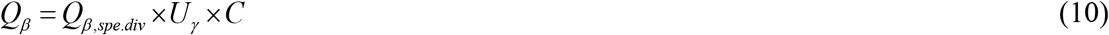

*Q*_*β,spe*.*div*_ and *U*_*γ*_ vary between 0 and 1; *C* is also positive but can be higher than 1 in case of FP clustering. In real case study, we expect that various values of *C* will be observed depending on the ecological processes that act in the study area. We expect that *U*_*γ*_ be low to moderate in most case studies as at a regional level, species from a restricted taxonomic group usually share part of their FP characteristics, although for trait-based and functional diversity this will depend on the number of traits considered and on the level of correlations between traits. The component that we expect to be often low is *Q*_*β,spe*.*div*_. Indeed, although abundance distribution within a local community is known to be asymmetric with a few species being the most abundant and many other species being rare, the strong abundance-driven clustering where a single species dominates in abundance in each community but the dominant species is different from one area to another is likely to be uncommon.

Overall the values of *Q*_*β*_ are thus expected to be low in real case study (as previously observed, e.g., by Hardy & Jost, 2008). However comparing *Q*_*β*_ / *Q*_*γ*_ between different regions will inform on the level of FP clustering in these regions. And comparing each component of *Q*_*β*_ between different regions can be even more informative as it will reveal how regions differ in terms of relative FP clustering and γ FP uniqueness.

The expression of a link between *D*_*β*_ and *D*_*β*,*spe*.*div*_ has to take a different form than that of the link between *Q*_*β*_ and *Q*_*β*,*spe*.*div*_ . This is because, while *Q*_*β*_ and *Q*_*γ*_ are expressed in the same unit, *D*_*β*_ is not commensurate with *D*_*γ*_ : *D*_*β*_ is expressed in terms of effective number of areas while *D*_*γ*_ expresses an effective number of species. A simple way to depict the relationship between *D*_*β*_ and *D*_*β*,*spe*.*div*_ is thus to consider *D*_*β*_ = *U*_*R*_ × *D*_*β*,*spe*.*div*_, with:

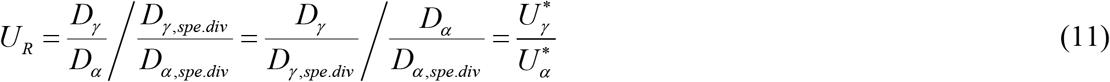

*U*_*R*_ is the ratio of γ (regional) uniqueness 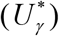 to α (local) uniqueness 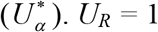 both when areas have identical compositions and when they are maximally dissimilar. *U*_*R*_ is high (>1) in case of high γ uniqueness (high 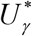) and low α uniqueness 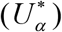, i.e. high α redundancy. High values of *U*_*R*_ are achieved notably when the local FP diversity is low, local uniqueness is also low in part of the areas (high species diversity, but species are similar, having similar traits or close phylogenetic positions), and the species which are the most distinct in the species pool are the most ubiquitous, which increases their relative abundance at the γ level compared to the α level. In contrast, *U*_*R*_ is low (<1) when each species in a given area tends to have sibling species (with similar FP characteristics) in the other areas (an example of these two contrasting cases will be given in the Results section).

### Case studies

We considered, as an illustration, simulated theoretical datasets based on a pool of 100 species characterized by a single quantitative, discrete trait. To obtain distances between species bounded in 0 and 1, we first standardized the trait by dividing it by the range of variation (maximum-minimum) and defined the dissimilarity between two species has the absolute difference in their standardized trait value. Note that only *D*_*β*_ requires the use of bounded distances but for homogeneity we apply also the standardization when using *Q*_*β*_ . Let *y*_*i*_ be the raw trait value for species *i*; and *x*_*i*_ = *y*_*i*_ / (max_*j*_(*x*_*j*_)-min_*j*_(*x*_*j*_)) be its standardized trait value.

We simulated functional diversity in 5 areas. We considered a reference species pool with 100 species with raw trait values (the *y*_*i*_’s) equal to 1 to 100 with a step of 1. The number of individuals in the species pool was not fixed (considered infinite); however, we considered species as evenly abundant in the species pool. Then we considered that 250 individuals colonized each area from the species pool.

- In the first scenario (randomness), the 250 individuals were randomly drawn from the species pool.
- In the second scenario (clustering), each area was associated with an optimal trait value to simulate functional clustering. The probability that an individual of species *i* entered the area *m* was inversely linked to the difference between its trait value and the optimal trait value of area *m* by the following formula: 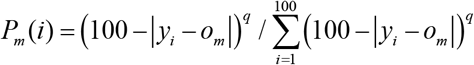, where *o*_*m*_ is the optimum trait value in area *m* and *q* is a positive exponent that controls the strength of the filtering, with *q* = 1 the minimum considered strength. We considered four values for *q*: 1, 3, 9 and 81. Optimal trait values for the five areas were defined as *o*_1_ = 5, *o*_2_ = 28, *o*_3_ = 50, *o*_4_ = 72, and *o*_5_ = 95, respectively. We also considered the extreme case where only the species that had the optimal trait value was present in the area (i.e., *q* → ∞).
- In the third scenario (overdispersion), an individual was first randomly drawn from a species pool to enter an area. New individuals were iteratively added to the area with the following constraint: the probability that a given individual was selected to enter the area is linked to the average absolute difference between its trait value and the trait values in the set Σ of species already present in the area by the following formula: 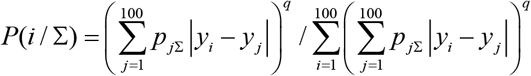, where *p*_*jΣ*_ is the relative abundance of species *j* in the set Σ, and *q* is a positive exponent that controls the strength of the overdispersion, with *q* = 1 the minimum considered strength. We considered four values for *q*: 1, 3, 9 and 81.
- In the fourth scenario (outliers), the areas were submitted to different pressures: one area was composed of the first 50 species of the species pool (with trait values between 1 and 50) and the four others were composed of two atypical species, which do not belong to the reference species pool, with trait values 499 and 500, respectively; abundances were random with the only constraint that each area was composed of 250 individuals.
- In the fifth scenario (siblings), 5 species were first randomly drawn from the species pool (except species with the extreme trait values 1 and 100). These 5 species constituted the first area. Then the composition of the other areas was also based on the set of 5 selected species, except that each of these 5 species could be randomly replaced by one of its closest neighbor (a species with a trait value of *y* has two closest species with trait values *y*-1 and *y*+1, respectively). Species with extreme trait values were excluded in the first area as these only have one closest neighbor. As in the fourth scenario, the abundances of the species in each area were randomly defined with the only constraint that the area was composed of 250 individuals.

For each scenario, 1000 sets of 5 areas were simulated and the components *Q*_*β*_, *U*_*γ*_, *C, Q*_*β,spe*.*div*_, and *D*_*β*_, *U*_*R*_, *U* and *D*_*β,spe*.*div*_ were computed on each set.

## RESULTS

Scenarios 1 to 3 allowed us to illustrate the behavior of the *Q*_*β*_ index and its components *C, U* and *Q*_*β,spe*.*div*_. The functional β diversity (*Q*_*β*_) increased with the strength of the clustering (Fig. 1). Species β diversity (*Q*_*β,spe*.*div*_) increased more slowly than functional β diversity with the strength of the clustering but it reached higher values (Fig. 1). The *γ* uniqueness (*U*_*γ*_) increased as either the strength of the overdispersion or that of the clustering increased (Fig. 1). The *C* index increased from high overdispersion, through the random scenario, to peak at moderate clustering and then decreased for high clustering as functional clustering could not be distinguished from species clustering (Fig. 1). When the species were randomly distributed into the areas, the value of the *C* index, as expected, approached unity (Fig. 1).

**FIGURE 1.**
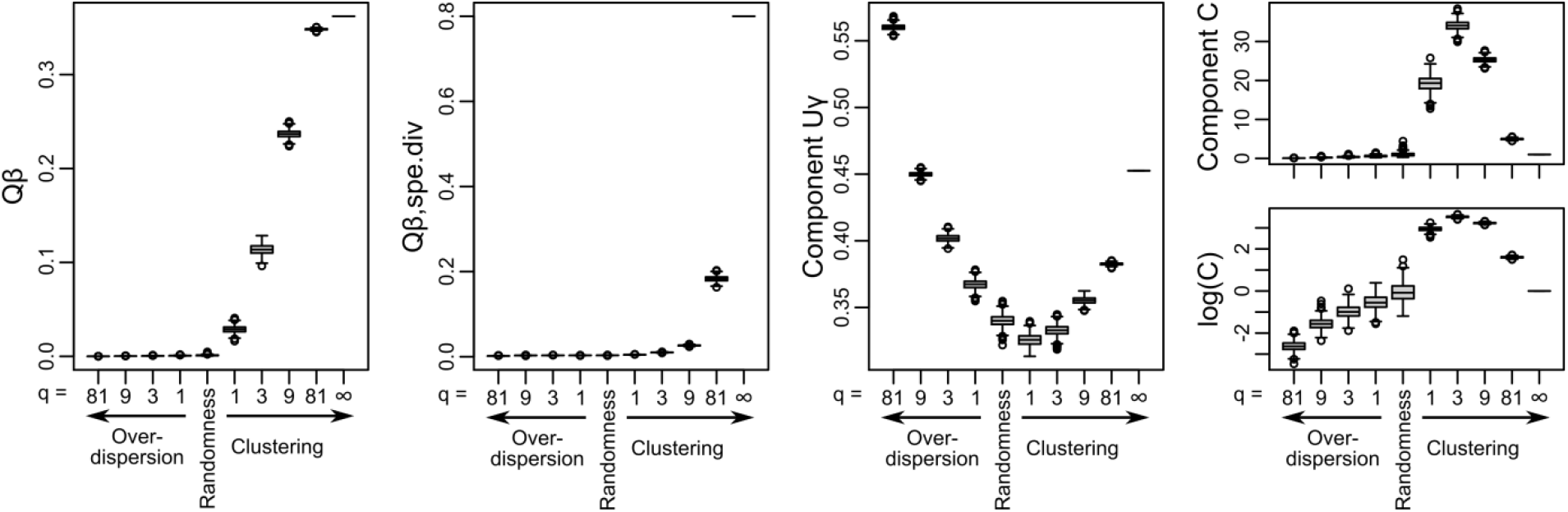
Level of functional β diversity, as measured by *Q*_*β*_, and its components obtained for each scenario of the case study. From left to right and top-right to bottom-right: functional β diversity (*Q*_*β*_), species β diversity (*Q*_*β, spe*.*div*_), γ functional uniqueness (*U*_*γ*_), functional clustering (*C*), log-transformed component of functional clustering (log(*C*)). Values are summarized by a box plot for each simulation scenario from overdispersion with decreasing strength (Scenario 3 in section *Case Studies*, with parameter *q* = 81, 9, 3, 1), through random simulation (Scenario 1 in section *Case Studies*), to clustering with increasing strength (Scenario 2 in section *Case Studies*, with parameter *q* = 1, 3, 9, 81, and ∞).

Comparing scenarios 1, 4 and 5 illustrated the behavior of the uniqueness ratio (*U*_*R*_) index (ratio of *D*_*β*_ to *D*_*β,spe*.*div*_, also equal to the ratio of *γ* uniqueness to *α* uniqueness). The *U*_*R*_ values increased from scenario 5 where each species in a given area tended to have sibling species (with similar FP characteristics) in the other areas to scenario 4 where an area has 50 species with trait values between 1 and 50 and the four other areas have two comparatively atypical species with trait values 499 and 500 (Fig. 2a). Scenario 1 (random distribution of species within areas) led to intermediate *U*_*R*_ values (Fig. 2a). Note that the values of *U*_*R*_ tend to unity under the random scenario as the number of individuals per area increases (Appendix S1).

**FIGURE 2.**
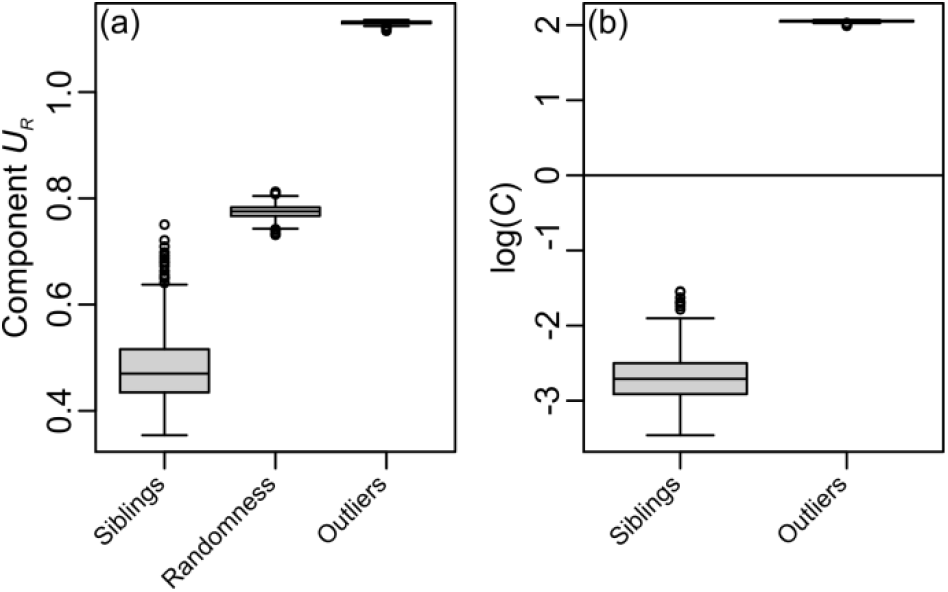
Distribution of components *U*_*R*_ (a) and (log-transformed) *C* (b) over simulations summarized by box plots. Box plots are given for three scenarios: Siblings (Scenario 5 in section *Case Studies*), Randomness (Scenario 1 in section *Case Studies*) and Outliers (Scenario 4 in section *Case Studies*).

Scenarios 1 to 3 revealed that, although as for *Q*_*β*_ functional *D*_*β*_ increases from overdispersion to clustering, species *D*_*β,spe*.*div*_ increases more rapidly than *D*_*β*_ with the strength of clustering. As a consequence, they also revealed that in general *U*_*R*_, the ratio of *D*_*β*_ to *D*_*β,spe*.*div*_, decreases from high overdispersion to high clustering (Fig. 3). This contradicts the results obtained with scenarios 4 and 5 (Fig. 2a). Scenarios 4 and 5 are indeed, respectively, particular cases of clustering and overdispersion, so that *C* took high (>>1) values under scenario 4 and low values (<<1) under scenario 5 (Fig. 2b). However, they represent special cases of clustering and overdispersion, respectively, that depart from the general tendency. When overdispersion is high then both *D*_*β*_ to *D*_*β,spe*.*div*_ approach unity so that *U*_*R*_ also approaches unity. We have to recall also that when areas are maximally dissimilar, then *D*_*β*_ = *D*_*β,spe*.*div*_ so that *U*_*R*_ = 1 and is thus, in that case, independent of the level of clustering within the areas. So *U*_*R*_ values close to unity are obtained in case of overdispersion or when areas are maximally dissimilar, independently of the level of clustering/overdispersion within the areas (Fig. 3).

**FIGURE 3.**
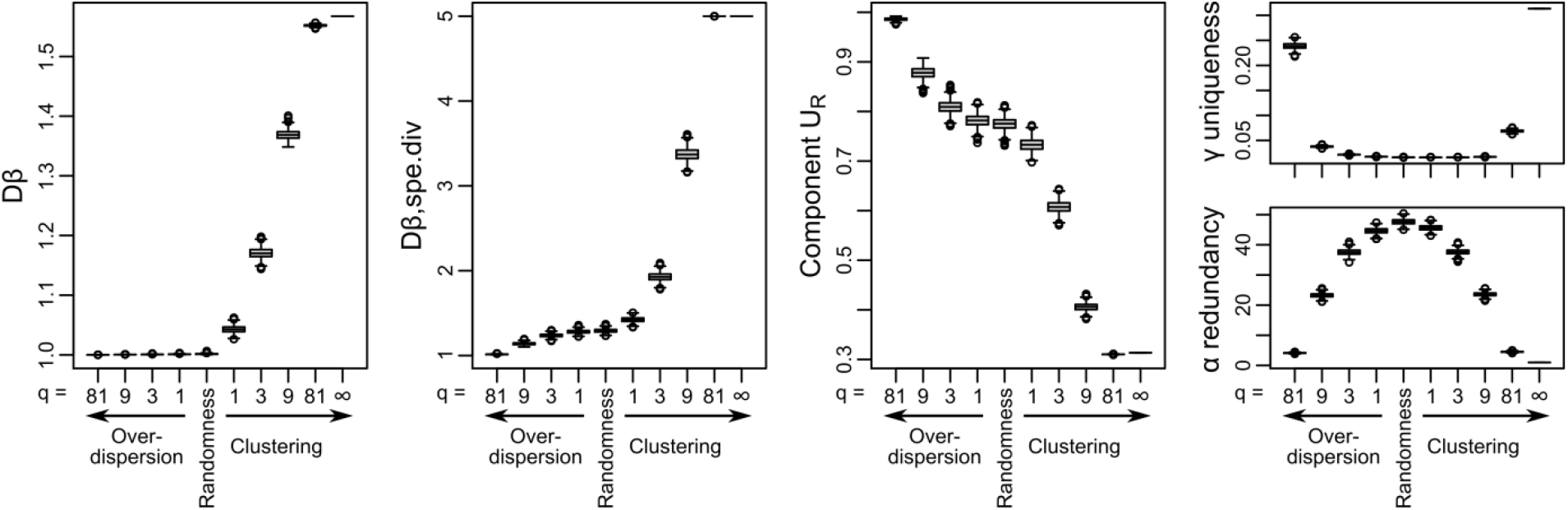
Level of functional β diversity, as measured by *D*_*β*_, and its components obtained for each scenario of the case study. Distributions, over simulations, are given for (from left to right and top-right to bottom-right) functional β diversity (*D*_*β*_), species β diversity (*D*_*β,spe*.*div*_), the ratio *U*_*R*_ of functional to species β diversity, functional uniqueness at the γ level 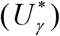, and functional redundancy at the α level 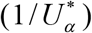 Values are summarized by a box plot for each simulation scenario from overdispersion with decreasing strength (Scenario 3 in section *Case Studies*, with parameter *q* = 81, 9, 3, 1), through random simulation (Scenario 1 in section *Case Studies*), to clustering with increasing strength (Scenario 2 in section *Case Studies*, with parameter *q* = 1, 3, 9, 81, and ∞).

## DISCUSSION

### Disentangling diversity indices into elementary components

Here we used a new way of partitioning FP β diversity into elementary components to reveal how FP β diversity depends on the concepts of clustering/overdispersion and redundancy/uniqueness. One of these elementary components is species β diversity. The partitioning we propose thus allows identifying when and why FP β diversity differs from species β diversity. Global evaluation of the ratio of FP β diversity to species β diversity has been done at different scales in several recent studies: between continents, islands, regions, or other geographic locations (e.g., Zhao et al., 2021); between biomes, ecosystems, successional stage, habitats, or areas within an habitat (e.g., Ricotta et al., 2020); or also between hosts for a parasite group (Krasnov et al., 2021).

Analyzing this ratio in terms of elementary components can help studying what insights on β diversity are gained by using trait-based, functional or phylogenetic information compared to more simply considering species abundance. Notably, at broad geographic scale, the combined effects of dispersal limitation, association with a biome, particular history of speciation and extinction, and niche conservatism may lead to decoupling of phylogenetic β diversity with species β diversity (Azevedo et al., 2021). At more local scales, the decoupling of FP β diversity with species β diversity may be due for example to disturbance that affects the level of habitat heterogeneity and habitat filtering (e.g., Rodrigues et al., 2017) or to modified patterns of interspecific competitions (e.g., Wong et al., 2019).

Species invasion driven by human activities, for example, can reduce functional β diversity while maintaining species β diversity (e.g., Wong et al., 2019). However the reasons why this is the case still have to be elucidated. Partitioning FP β diversity into elementary components could help. Wong et al. notably observed the removal of functionally unique species at the γ level in invaded communities. In that case the cumulative removal of the few unique species from each invaded community would yield a massive decrease in functional uniqueness at the γ level. γ uniqueness being one of the elementary component of both *Q*_*β*_ and *D*_*β*_ indices analyzed above, a decrease in functional β diversity in invaded communities, leading to a functional homogenization at the landscape scale, might thus result from the loss of the functionally unique species of communities. Assessing elementary components in FP β diversity could thus allow a finer description of non-random patterns of community change due to human activities (e.g., Uchida et al., 2018).

Several methodological studies have tried to partition FP diversity indices into elementary components to disentangle the parameters responsible for changes in FP diversity in light of various factors including spatial scale, environmental change, and disturbance (e.g., Patil & Taillie, 1982; Schmera et al., 2020; Shimatani, 2001). This is because different components may reveal different mechanisms and processes of community assembly. As regards indices of β diversity, previous studies that managed to partition β diversity into elementary components often expressed these components in terms of nestedness, richness difference (species gains/losses), turnover or replacement (Schmera et al., 2020). Such extraction of elementary components is now done frequently on species diversity (e.g., Schmera et al., 2020) and on FP diversity (e.g., Melo et al., 2014). However the influence of FP clustering versus overdispersion and those of FP redundancy versus uniqueness on FP β diversity have so far been far less studied.

### β diversity and clustering/overdispersion

Usually clustering/overdispersion patterns are not identified over a whole studied region but within each local area in reference to a large delimited region (e.g., Molina-Venegas et al., 2022; Ramm et al., 2018; Rocha et al., 2019; Yang et al., 2014). Their identification has so far required the use of null models and randomization of species occurrences or FP characteristics (e.g., Hardy & Senterre, 2007; Webb et al., 2002). In Webb et al. (2002) seminal paper, clustering is, among other approaches, identified in a given area if the average phylogenetic dissimilarity between two species in the area is lower than that expected by randomly assembling species in the area from a larger, regional species pool (NRI index). In contrast, overdispersion is identified if the average dissimilarity between two species in the area is higher than that expected by randomly assembling species in the local area from the regional species pool. The distribution of values obtained for the NRI index of clustering/overdispersion can then be summarized by analyzing their mean, standard deviation and/or quantiles to identify general trends (e.g., Luo et al., 2021; Rocha et al., 2019; Yang et al., 2014). Webb et al. (2002) used presence-absence data. Nevertheless, the *Q* index, which evaluates the average abundance-weighted dissimilarity within an area, was also found as one of the metrics the most efficient in the detection of clustering and overdispersion when it is associated with a relevant null model of random community assembly (Miller et al., 2017).

Here we showed that *Q*_*β*_, in the scale-based partitioning of *Q*, has an intrinsic component (*C*) that directly summarizes the global level of clustering over the studied region: *C* is low in case of an overall trend towards overdispersion and high in case of an overall trend towards clustering. Opting for the use of this component *C*, the detection of an overall trend of overdispersion or clustering does thus not require the involvement of null models. This is reassuring given the difficulty there is to identify best performing null models (Hardy, 2008; Miller et al., 2017), the additional methodological step they represent, and given the amount of calculation these null models require, particularly on large datasets.

Although index *C* could also be used with null models to test the significance of an observed clustering or overdispersion, it has thus the great advantage of evaluating levels of clustering versus overdispersion without the need for these null models. This can be useful in situations where the aim is to compare the level of clustering versus overdispersion of distinct sets of communities be they for example, different ecoregions (to reveal, e.g., the influence of habitat fragmentation, habitat structure or local climate in shaping functional clustering), continents (to reveal, e.g., geographic or climate related effects), or altitudinal strata (to reveal, e.g., an increase in functional clustering with elevation). Indeed in many studies, it is not so much the significance of a clustering or overdispersion in each community that is of interest but rather that of the trend of variation in clustering, such as the increase in clustering along a gradient (Miller et al., 2017).

By analyzing Rao’s quadratic entropy we showed that the switch from a dissimilarity-based framework (*Q*_*β*_) to an equivalent-number framework (*D*_*β*_) has consequences in terms of how clustering versus overdispersion is expected to influence FP β diversity relatively to species β diversity. *Q*_*β*_ is a measure of average between-communities dissimilarity, while *D*_*β*_ is an equivalent number of areas. In *Q*_*β*_, FP β diversity is expected to be low in case of high functional overdispersion. This is because low functional clustering means that, on average, two species drawn from two different areas are as functionally similar as two species drawn from the same area.

In our simulations, although we observed that *C* globally increased from functional overdispersion to functional clustering, we also observed that, as expected from a relative clustering index, *C* decreased with extremely high functional clustering. Indeed, *C* is an index of FP clustering standardized by species clustering. When the strength of functional clustering is extremely high, then less and less species are shared between areas, so that the functional clustering (when two species in an area are more functionally similar to each other than are two species one from an area and the other from another area) is confounded with species clustering (when areas do not share species and each area is characterized by the presence/dominance of a few species, whatever their traits). When the strength of the functional clustering becomes extremely high, it can thus hardly be distinguished from a species clustering. In the extreme scenario where each area contains only the species that has the optimal trait, *C* = 1, meaning that, with the only information provided to *C* formula, it becomes impossible to evaluate whether the clustering is due to the traits of the species or to something else.

With *D*_*β*_, the ratio *D*_*β*_ to *D*_*β,spe*.*div*_ generally decreases from high overdispersion to high clustering but there are exceptions when the overdispersion and clustering take particular forms, like in scenarios 4 and 5 of our simulations. In the general case, with high overdispersion, both *D*_*β*_ and *D*_*β,spe*.*div*_ tend to values approaching 0 (and the ratio tends to 1), while, with high clustering, the areas tend to not share species any more (i.e. *D*_*β,spe*.*div*_ tends to its maximum value) but still share some functional characteristics (i.e. *D*_*β*_ has moderate to high but not maximum values). In scenarios 4 and 5 we considered, either some areas had atypical trait values (reflecting their high uniqueness; scenario 4), or areas all had a large range of trait value but for each species in an area, there was a species in every other area whose trait value is close to its own (reflecting high redundancy between areas; scenario 5). Our analyses of *Q*_*β*_ and *D*_*β*_ thus revealed that both indices are influenced by FP clustering, although only *Q*_*β*_ is a simple, increasing function of it. Whether FP β diversity should increase with FP clustering is thus an open question that our analyses raise. The response to this question will likely depend on the objective of a study (e.g., Obj1, Obj2 versus Obj3 of the Introduction section).

### β diversity as a function of α and γ uniqueness

Comparing FP diversity patterns with species diversity patterns can help reveal community assembly processes (e.g., Pavoine & Bonsall, 2011; Larson et al., 2021). This is because high species diversity accompanied by relatively low FP diversity indicates FP redundancy and low species diversity compensated by relatively high FP diversity indicates FP uniqueness. Phylogenetic redundancy/uniqueness patterns can be signatures of evolutionary processes such as diversification and dispersal between regions. For example, at the regional level, phylogenetic redundancy may be due to recent speciation events or to particular species’ extinction events that affected old lineages leaving one recent clade. Inversely, phylogenetic uniqueness could be due to the colonization of an area by phylogenetically-distant species with different geographic origins, to a lack of radiation, to low speciation rate, or to frequent extinction events that pruned all clades (e.g., Qian & Deng, 2021). In complement, functional redundancy/uniqueness patterns can be signatures of ecological processes such as avoidance of niche overlap by competitive exclusion for functional uniqueness. In undisturbed environments with unlimited resources hampering competition, high functional redundancy can be associated with environmental filtering that drives the co-occurrence of a large set of specialist species with similar traits adapted to local favorable environmental conditions, leading to high species diversity but low functional diversity. In rich systems, α and *γ* functional redundancy have been associated with resilience and stability to disturbance (Biggs et al., 2020) and to the ability of a species set to fulfill a diverse set of ecosystem functions and services (e.g., Mori et al., 2016). Indeed, redundant species provide biological buffering capacity that leads to stable community functions. In case of high functional redundancy, one could indeed expect that the extinction of a redundant species can be compensated by an extant functionally-similar species (e.g., Johnson et al., 1996).

However for this statement to be correct, both functional diversity and species diversity have to be high (e.g., Aguirre-Gutiérrez et al., 2022) and variation in functional redundancy have to result from variation in species diversity only (biodiversity-ecosystem functioning insurance hypothesis, Yachi & Loreau, 1999). For example, in highly-diverse ecosystems, fluctuations in species diversity during seasonal disturbances may not be accompanied by similar fluctuations in functional diversity if the decrease in abundance for certain species is compensated by the increase in abundance of functionally redundant species (e.g., Wang et al., 2023). In that case, the accumulation of functional redundancy during favorable season allow a stabilization of functional diversity during the least favorable seasons. If, in contrast, an increase of functional redundancy results from biodiversity loss with more drastic loss in functional diversity than the loss of species diversity as might happen when functionally unique species are lost first under disturbance, then functional redundancy could be associated with irreplaceable loss of ecosystem functions (e.g., Flynn et al., 2009).

Here we showed that two widely-used indices of β diversity are related to measures of FP uniqueness. By our partitioning of FP β diversity indices, we highlighted that the ratio of FP β diversity to species β diversity is itself dependent on FP uniqueness at the γ level for *Q*_*β*_ and equal to the ratio of γ uniqueness to α uniqueness for *D*_*β*_. The level of FP redundancy might indeed differ between spatial scales so that γ redundancy is likely to differ from α redundancy (e.g., Lamothe et al., 2018). Variations in FP redundancy at the α (for *D*_*β*_) and γ levels (for *D*_*β*_ and *Q*_*β*_) may thus drive changes in the ratio of FP β diversity to species β diversity.

Although redundancy has still rarely been discussed in a context of β diversity, we show here that this concept implicitly enters in the definition of well-used β diversity indices. The relationship between local redundancy, regional redundancy and β diversity has thus to be further explored in a variety of ecosystems in lights of ecological and evolutionary processes. Finally, as FP redundancy and uniqueness are known to be dependent on the number and identity of functions considered (e.g., Mori et al., 2016), this further supports the fact that trait selection is important in the measurement of functional β diversity. For a direct link with ecosystem functions, functional traits have to be selected as the organismal traits that influence one or more aspects of the functioning of an ecosystem (Tilman, 2001). The number of ecosystem functions considered, the number of traits considered and the correlations between them might affect the level of redundancy observed and thus the level of functional β diversity according to both indices we considered but especially according to *Q*_*β*_ that only depends on the absolute level of γ redundancy, while in *D*_*β*_ it is evaluated relatively to the level of α redundancy. Large levels of α, β, γ functional diversity in reference to species diversity are more likely to be observed with the consideration of a large number of uncorrelated traits than if a few correlated traits are selected.

### Conservation studies versus community ecology studies

The measurement of biodiversity cannot lead to a unique mathematical formula (Hoffman a&nd Hoffman, 2008). However, biodiversity indices must be backed by sound argument (or, more merely, must be reasonable; Hoffman & Hoffman, 2008) regarding the objective of a study. A biodiversity index actually is an index, not of all biodiversity, but of some specific aspect of biodiversity that has to be clearly specified and understood to facilitate the interpretation of its values. Partitioning an index into elementary components helps in identifying the specific aspects of biodiversity that it expresses (e.g., Patil & Taillie, 1982; Shimatani, 2001). For β diversity, the partitioning we introduced here, where FP β diversity is expressed in terms of species β diversity among other components, complements the approaches that partition FP β diversity into gains/losses, nestedness, or turnover/replacement components. Such partitions into elementary components are useful to better identify which aspects of FP β diversity each index expresses and to search for the mechanisms that have shaped biodiversity patterns in a study area.

Our partitioning of the related indices *Q*_*β*_ and *D*_*β*_ shows that *D*_*β*_ relies on a traditional view of biodiversity in conservation biology that is based on counts (safeguarding the highest possible number of species, and the highest possible number of species communities) and that *Q*_*β*_ can be apportioned into components usually used in community ecology to detect underlying ecological processes (FP clustering and FP redundancy). Rao’s apportionment of quadratic entropy models community systems considering that FP β diversity increases with species β diversity, FP clustering at the α level (i.e., within areas) and uniqueness at the γ level (i.e., over all combined areas). In the introduction, we indicated three frequent objectives for measuring biodiversity: “Obj1” saving biodiversity for its intrinsic value, “Obj2” maintaining ecosystem functioning and services, and “Obj3” understanding the processes underlying biodiversity patterns. Clearly, *D*_*β*_ definition is more intuitive for communication to a large public in the context of Obj1 as *D*_*β*_ is an effective number of communities. Indeed, without a thorough explanation of *Q*_*β*_ definition in link with the concepts of clustering and overdispersion, using it for communication to policy making is taking the risk of misinterpretation of its value.

But knowledge of the processes involved in biodiversity dynamics is important for long-term conservation strategies. This is where research in community ecology and in conservation biology come together: reaching objective Obj1 also requires the implementation of objectives Obj2 and Obj3. The redundancy components that both *Q*_*β*_ and *D*_*β*_ contain are particularly informative in the context of Obj2. As for Obj3, a critical difference between *Q*_*β*_ and *D*_*β*_ is that in addition to species β diversity and FP γ uniqueness, *Q*_*β*_ increases with FP clustering while *D*_*β*_ increases with α FP redundancy. Clustering and α redundancy are related but different concepts. While clustering implies that two species co-occurring in a common site tend to have more similar FP characteristics than two species from different sites, α redundancy only requires that co-occurring species tend to have similar FP characteristics. As shown in our simulations, there exist scenarios where the ratio of γ uniqueness to α uniqueness does not reflect clustering/overdispersion.

## CONCLUSION

We have shown here that opting for the additive or the multiplicative decomposition of the quadratic entropy modifies how FP β diversity relates to species β diversity. We also underlined that it modifies the definition of species β diversity itself: one, for the additive decomposition, based on the deviation from a case of complete species clustering where there are as many studied areas as species and areas do not share species; and the other, for the multiplicative decomposition, based on the excess in abundance evenness among species at the γ level compared to that at the α level. More generally, ecologists have to be fully aware that currently there is no single, consensual definition of FP β diversity so that opting for an index over another is not innocuous and has important consequences on the aspect of β diversity that they are measuring.

The relationship between species diversity and FP diversity that we analyzed here is poorly documented at all scales and particularly at the β level. This comparison is however important both to evaluate the processes that shape community assembly and also to evaluate the resilience and stability abilities of an ecosystem at a landscape level. An alternative way to evaluate how FP β diversity differs from species β diversity would be to compare the observed value of FP β diversity with null expectations based on species β diversity (e.g., Azevedo et al., 2020). However the decompositions of *Q*_*β*_ and *D*_*β*_ we propose here allow identifying more precisely why FP β diversity differs from species β diversity: For *Q*_*β*_, is it in terms of γ uniqueness or in terms of clustering/overdispersion? For *D*_*β*_, is it because of particularly close, or inversely, particularly dissimilar values of γ uniqueness and α uniqueness? From this study that focuses on two FP β diversity indices, we call for thorough analyses of existing indices of FP β diversity in terms of how much they are driven by patterns of clustering/overdispersion and/or redundancy/uniqueness. This will ensure that the relevance of an index in light of the specified, selected objective of a study has been checked and that the values taken by the index on field data can be interpreted with the whole information they convey. We also call for further studies on whether/when/why functional and/or phylogenetic β diversity should increase with FP clustering.

## AUTHOR CONTRIBUTIONS

S.P. conceived and designed the study; S.P. conducted the analyses; S.P. and C.R. analyzed the results and contributed to write the paper.

## REFERENCES

Aguirre-Gutiérrez, J., E. Berenguer, I. Oliveras Menor, D. Bauman, J. J. Corral-Rivas, Nava-M. G. Miranda, S. Both, et al. 2022. Functional susceptibility of tropical forests to climate change. Nature Ecology and Evolution 6: 878–889. 10.1038/s41559-022-01747-6

Albert, J. S., G. Destouni, S. M. Duke-Sylvester, A. E., Magurran, T. Oberdorff, R. E. Reis, K. O. Winemiller, and W. J. Ripple. 2021. Scientists’ warning to humanity on the freshwater biodiversity crisis. Ambio 50: 85–94. 10.1007/s13280-020-01318-8

Azevedo, J. A. R., C. C. Nogueira, A. Antonelli, and S. Faurby. 2020. Contrasting patterns of phylogenetic turnover in amphibians and reptiles are driven by environment and geography in Neotropical savannas. Journal of Biogeography 48: 2008–2021. 10.1111/jbi.14131

Bañares-de-Dios, G., M. J. Macía, Í. Granzow-de la Cerda, I. Arnelas, G. Martins de Carvalho, C. I. Espinosa, N. Salinas, et al. 2020. Linking patterns and processes of tree community assembly across spatial scales in tropical montane forests. Ecology 101: e03058. 10.1002/ecy.3058

Biggs, C. R., L. A. Yeager, D. G. Bolser, C. Bonsell, A. M. Dichiera, Z. Hou, S. R. Keyser, et al. 2020. Does functional redundancy affect ecological stability and resilience? A review and meta-analysis. Ecosphere 11(7): e03184. 10.1002/ecs2.3184

Borges, P. A., F. Rigal, A. Ros-Prieto, and P. Cardoso. 2020. Increase of insular exotic arthropod diversity is a fundamental dimension of the current biodiversity crisis. Insect Conservation and Diversity 13: 508–518. 10.1111/icad.12431

Carta, A., L. Peruzzi, and S. Ramírez-Barahona. 2022. A global phylogenetic regionalization of vascular plants reveals a deep split between Gondwanan and Laurasian biotas. New Phytologist 233: 1494–1504. 10.1111/nph.17844

Cavender-Bares, J., D. D. Ackerly, D. A. Baum, and F. A. Bazzaz. 2004. Phylogenetic overdispersion in Floridian oak communities. The American Naturalist 163: 823–843. 10.1086/386375

Chao, A., C. H. Chiu, and T. C. Hsieh. 2012. Proposing a resolution to debates on diversity partitioning. Ecology 93: 2037–2051. 10.1890/11-1817.1

Chen, K., A. R. Rajper, R. M. Hughes, J. R. Olson, H. Wei, and B. Wang. 2019. Incorporating functional traits to enhance multimetric index performance and assess land use gradients. Science of The Total Environment 691: 1005–1015. 10.1016/j.scitotenv.2019.07.047

Chiu, C. H., L. Jost, and A. Chao. 2014. Phylogenetic beta diversity, similarity, and differentiation measures based on Hill numbers. Ecological Monographs 84: 21–44. 10.1890/12-0960.1

Cooper, N., J. Rodríguez, and A. Purvis. 2008. A common tendency for phylogenetic overdispersion in mammalian assemblages. Proceedings of the Royal Society B: Biological Sciences 275: 2031–2037. 10.1098/rspb.2008.0420

Davis, M., S. Faurby, and J. C. Svenning. 2018. Mammal diversity will take millions of years to recover from the current biodiversity crisis. Proceedings of the National Academy of Sciences of the USA 115: 11262–11267. 10.1073/pnas.180490611

Flynn, D. F., M. Gogol-Prokurat, T. Nogeire, N. Molinari, B. T. Richers, B. B. Lin, N. Simpson, et al. 2009. Loss of functional diversity under land use intensification across multiple taxa. Ecology Letters 12: 22–33. 10.1111/j.1461-0248.2008.01255.x

Gallien, L. 2017. Intransitive competition and its effects on community functional diversity. Oikos 126: 615–623. 10.1111/oik.04033

González, C., R. Macip-Ríos, and I. Suazo-Ortuño. 2022. Phylogenetic structure and diversity among herpetofaunal communities along a successional gradient of a tropical dry forest in Mexico. Perspectives in Ecology and Conservation 20: 249–255. 10.1016/j.pecon.2022.05.004

Gower, J. C., and P. Legendre. 1986. Metric and Euclidean properties of dissimilarity coefficients. Journal of Classification 3: 5–48. 10.1007/bf01896809

Hardy, O. J. 2008. Testing the spatial phylogenetic structure of local communities: statistical performances of different null models and test statistics on a locally neutral community. Journal of Ecology 96: 914–926. 10.1111/j.1365-2745.2008.01421.x

Hardy, O. J., and L. Jost. 2008. Interpreting and estimating measures of community phylogenetic structuring. Journal of Ecology 96: 849–852. 10.1111/j.1365-2745.2008.01423.x

Hardy, O. J., and B. Senterre. 2007. Characterizing the phylogenetic structure of communities by an additive partitioning of phylogenetic diversity. Journal of Ecology 95: 493–506. 10.1111/j.1365-2745.2007.01222.x

Hoffmann, S., and A. Hoffmann. 2008. Is there a “true” diversity? Ecological Economics 65: 213–215. 10.1016/j.ecolecon.2008.01.009

Johnson, K. H., K. A. Vogt, H. J. Clark, O. J. Schmitz, and D. J. Vogt. 1996. Biodiversity and the productivity and stability of ecosystems. Trends in Ecology & Evolution 11(9): 372–377. 10.1016/0169-5347(96)10040-9

Jost, L. 2006. Entropy and diversity. Oikos 113: 363–375. 10.1111/j.2006.0030-1299.14714.x

Jurasinski, G., V. Retzer, and C. Beierkuhnlein. 2009. Inventory, differentiation, and proportional diversity: a consistent terminology for quantifying species diversity. Oecologia 159, 15–26. 10.1007/s00442-008-1190-z

Krasnov, B. R., A. Spickett, K. Junker, L. van der Mercht, and S. Matthee. 2021. Functional and phylogenetic uniqueness of helminth and flea assemblages of two South African rodents. International Journal for Parasitology 51: 865–876. 10.1016/j.ijpara.2021.02.003

Lamothe, K. A., K. M. Alofs, D. A. Jackson, and K. M. Somers. 2018. Functional diversity and redundancy of freshwater fish communities across biogeographic and environmental gradients. Diversity and Distributions 24: 1612–1626. 10.1111/ddi.12812

Lande, R. 1996. Statistics and partitioning of species diversity, and similarity among multiple communities. Oikos 76: 5–13. 10.2307/3545743

Lanier, H. C., D. L. Edwards, and L. L. Knowles. 2013. Phylogenetic structure of vertebrate communities across the Australian arid zone. Journal of Biogeography 40: 1059–1070. 10.1111/jbi.12077

Larson, E. I., N. L. Poff, W. C. Funk, R. A. Harrington, B. C. Kondratieff, S. G. Morton, and A. S. Flecker. 2021. A unifying framework for analyzing temporal changes in functional and taxonomic diversity along disturbance gradients. Ecology 102: e03503. 10.1002/ecy.3503

Lavorel S., and É. Garnier. 2002. Predicting changes in community composition and ecosystem functioning from plant traits: revisiting the Holy Grail. Functional ecology 16: 545–556. 10.1046/j.1365-2435.2002.00664.x

Li, S. P., M. W. Cadotte, S. J. Meiners, Z. S. Hua, L. Jiang, and W. S. Shu. 2015. Species colonisation, not competitive exclusion, drives community overdispersion over long-term succession. Ecology Letters 18: 964–973. 10.1111/ele.12476

Luo, W., R. Lan, D. Chen, B. Zhang, N. Xi, Y. Li, S. Fang, et al. 2021. Limiting similarity shapes the functional and phylogenetic structure of root neighborhoods in a subtropical forest. New Phytologist 229: 1078–1090. 10.1111/nph.16920

Mayfield, M. M., and J. M. Levine. 2010. Opposing effects of competitive exclusion on the phylogenetic structure of communities. Ecology letters 13: 1085–1093. 10.1111/j.1461-0248.2010.01509.x

Melo, A. S., M. V. Cianciaruso, and M. Almeida-Neto. 2014. treeNODF: nestedness to phylogenetic, functional and other tree-based diversity metrics. Methods in Ecology and Evolution 5: 563–572. 10.1111/2041-210X.12185

Miller, E. T., D. R. Farine, and C. H. Trisos. 2017. Phylogenetic community structure metrics and null models: a review with new methods and software. Ecography 40: 461–477. 10.1111/ecog.02070

Molina-Venegas, R., G. Ottaviani, G. Campetella, R. Canullo, and S. Chelli. 2022. Biogeographic deconstruction of phylogenetic and functional diversity provides insights into the formation of regional assemblages. Ecography 2022: e06140. 10.1111/ecog.06140

Mori, A. S., F. Isbell, S. Fujii, K. Makoto, S. Matsuoka, and T. Osono. 2016. Low multifunctional redundancy of soil fungal diversity at multiple scales. Ecology letters 19(3): 249–259. 10.1111/ele.12560

Pasari, J. R., T. Levi, E. S. Zavaleta, and D. Tilman. 2013. Several scales of biodiversity affect ecosystem multifunctionality. Proceedings of the National Academy of Sciences of the USA 110: 10219–10222. 10.1073/pnas.1220333110

Patil, G. P., and C. Taillie. 1982. Diversity as a concept and its measurement. Journal of the American Statistical Association 77: 548–561. 10.2307/2287709

Pavoine, S., and M. Bonsall. 2011. Measuring biodiversity to explain community assembly: a unified approach. Biological Reviews 86, 792–812. 10.1111/j.1469-185X.2010.00171.x

Qian, H., and T. Deng. 2021. Geographic patterns and climate correlates of the deviation between phylogenetic and taxonomic diversity for angiosperms in China. Biological Conservation 262: 109291. 10.1016/j.biocon.2021.109291

Ramm, T., J. L. Cantalapiedra, P. Wagner, J. Penner, M. O. Rödel, and J. Müller. 2018. Divergent trends in functional and phylogenetic structure in reptile communities across Africa. Nature communications 9: 4697. 10.1038/s41467-018-07107-y

Rao, C. R. 1982. Diversity and dissimilarity coefficients: a unified approach. Theoretical Population Biology 21: 24–43. 10.1016/0040-5809(82)90004-1

Ren, Z., S. G. Baer, L. C. Johnson, M. B. Galliart, L. R. Wilson, and D. J. Gibson. 2023. The role of dominant prairie species ecotypes on plant diversity patterns of restored grasslands across a rainfall gradient in the US Great Plains. Applied Vegetation Science 26: e12725. 10.1111/avsc.12725

Ricotta, C. 2010. On beta diversity decomposition: Trouble shared is not trouble halved. Ecology 91: 1981–1983. 10.1890/09-0126.1

Ricotta, C., G. Bacaro, and S. Pavoine. 2015. A cautionary note on some phylogenetic dissimilarity measures. Journal of Plant Ecology 8: 12–16. 10.1093/jpe/rtu008

Ricotta, C., F. Laroche, L. Szeidl, and S. Pavoine. 2020. From alpha to beta functional and phylogenetic redundancy. Methods in Ecology and Evolution 11: 487–493. 10.1111/2041-210X.13353

Ricotta C., and L. Szeidl. 2009. Diversity partitioning of Rao’s quadratic entropy. Theoretical Population Biology 76: 299–302. 10.1016/j.tpb.2009.10.001

Rocha, J., R. E. Laps, C. G. Machado, and S. Campiolo. 2019. The conservation value of cacao agroforestry for bird functional diversity in tropical agricultural landscapes. Ecology and Evolution 9: 7903–7913. 10.1002/ece3.5021

Rodrigues, L. F., P. Cavalin, L. C. Franci, R. A. Bonaldi, V. Ariati, A. A. Padial, and M. C. Marques. 2017. Recurrent landslides affect the functional beta diversity of a megadiverse tropical forest. Plant Ecology & Diversity 10(5-6): 483–493. 10.1080/17550874.2018.1434568

Schmera, D., J. Podani, and P. Legendre. 2020. What do beta diversity components reveal from presence-absence community data? Let us connect every indicator to an indicandum! Ecological Indicators 117: 106540. 10.1016/j.ecolind.2020.106540

Shimatani, K. 2001. On the measurement of species diversity incorporating species differences. Oikos 93: 135–147.

Tilman, D. 2001. “Functional diversity”. In Encyclopedia of biodiversity,Vol. 3, edited by S. A. Levin, 109–120. San Diego, California, USA: Academic Press.

Tuomisto, H. 2010a. A diversity of beta diversities: straightening up a concept gone awry. Part 1. Defining beta diversity as a function of alpha and gamma diversity. Ecography 33: 2–22. 10.1111/j.1600-0587.2009.05880.x

Tuomisto, H. 2010b. A diversity of beta diversities: straightening up a concept gone awry. Part 2. Quantifying beta diversity and related phenomena. Ecography 33: 23–45. 10.1111/j.1600-0587.2009.06148.x

Uchida, K., M. Hiraiwa, and M. W. Cadotte. 2018. Non-random loss of phylogenetically distinct rare species degrades phylogenetic diversity in semi-natural grasslands. Journal of Applied Ecology 56: 1419–1428. 10.1111/1365-2664.13386

Veech, J. A., and T. O. Crist. 2010. Diversity partitioning without statistical independence of alpha and beta. Ecology 91: 1964–1969. 10.1890/08-1727.1

Wang, L., J. Li, L. Tan, and B.-P. Han. 2023. Seasonal patterns of functional alpha and beta redundancies of macroinvertebrates in a disturbed (sub)tropical river. Ecological Indicators 146: 109777. 10.1016/j.ecolind.2022.109777

Webb, C. O., D. D. Ackerly, and S. W. Kembel. 2008. Phylocom: software for the analysis of phylogenetic community structure and trait evolution. Bioinformatics 24: 2098–2100. 10.1093/bioinformatics/btn358

Webb, C. O., D. D. Ackerly, M. A. McPeek, and M. J. Donoghue. 2002. Phylogenies and community ecology. Annual review of ecology and systematics 33: 475–505. 10.1146/annurev.ecolsys.33.010802.150448

Weiher, E., and P. A. Keddy. 1995. Assembly rules, null models, and trait dispersion: new questions from old patterns. Oikos 74, 159–164. 10.2307/3545686

Whittaker, R. H. 1972. Evolution and measurement of species diversity. TAXON 21: 213–251. 10.2307/1218190

Wong, M. K. L., B. Guénard, and O. T. Lewis. 2019. The cryptic impacts of invasion: functional homogenization of tropical ant communities by invasive fire ants. Oikos 129: 585–597. 10.1111/oik.06870

Yachi, S., and M. Loreau. 1999. Biodiversity and ecosystem productivity in a fluctuating environment: The insurance hypothesis. Proceedings of the National Academy of Sciences of the USA 96: 1463–1468. 10.1073/pnas.96.4.1463

Yang, J., G. Zhang, X. Ci, N. G. Swenson, M. Cao, L. Sha, J. Li, et al. 2014. Functional and phylogenetic assembly in a Chinese tropical tree community across size classes, spatial scales and habitats. Functional Ecology 28: 520–529. 10.1111/1365-2435.12176

Zhao, Y., N. J. Sanders, J. Liu, T. Jin, H. Zhou, R. Lu, P. Ding, and X. Si. 2021. β diversity among ant communities on fragmented habitat islands: the roles of species trait, phylogeny and abundance. Ecography 44: 1568–1578. 10.1111/ecog.05723

