## Appendix S1 for "Functional and phylogenetic β diversities and their link with clustering/overdispersion and uniqueness/redundancy"

**Appendix S1. Supplementary information on Methods**

Sandrine Pavoine & Carlo Ricotta "Functional and phylogenetic β diversities and their link with clustering/overdispersion and uniqueness/redundancy"

### S1. Detailed equations for the apportionment of quadratic entropy (Q)

The value of the quadratic entropy applied to area *m* is

$$Q(\mathbf{p}_{im},\mathbf{D}) = \sum_{i=1}^{S_\gamma}\sum_{j=1}^{S_\gamma} p_{im}p_{jm}d_{ij} \quad \text{(S1)}$$

Rao (1982) defined the diversity within areas (α diversity) as

$$Q_\alpha = \sum_{m=1}^{M} w_m \sum_{i=1}^{S_\gamma}\sum_{j=1}^{S_\gamma} p_{im}p_{jm}d_{ij} \quad \text{(S2)}$$

the global diversity over all areas (γ diversity) as

$$Q_\gamma = Q\left(\sum\nolimits_{m=1}^{M} w_m\mathbf{p}_{im},\mathbf{D}\right) = \sum_{i=1}^{S_\gamma}\sum_{j=1}^{S_\gamma}\left(\sum_{m=1}^{M} w_m p_{im}\right)\left(\sum_{m=1}^{M} w_m p_{jm}\right)d_{ij} \quad \text{(S3)}$$

and the dissimilarity coefficient (DISC) between areas (β diversity) as

$$Q_\beta = Q_\gamma - Q_\alpha = \sum_{m=1}^{M} w_m \sum_{n=1}^{M} w_n \left(\sum_{i=1}^{S_\gamma}\sum_{j=1}^{S_\gamma} p_{im}p_{jn}d_{ij} - \frac{1}{2}\sum_{i=1}^{S_\gamma}\sum_{j=1}^{S_\gamma} p_{im}p_{jm}d_{ij} - \frac{1}{2}\sum_{i=1}^{S_\gamma}\sum_{j=1}^{S_\gamma} p_{in}p_{jn}d_{ij}\right) \quad \text{(S4)}$$

### S2. Interpretation of $Q_{\beta,spe.ric}$

Consider that FP dissmilarities ($d_{ij}$s) are bounded in [0, 1] and that $d_{ij}$ = 1 indicates that species *i* and *j* are maximally dissimilar. When species are maximally dissimilar and have similar abundances within each area, then in any area *m*, the relative abundance of any present species is $1/S_m$ with $S_m$ the species richness (number of species) in area *m* and the relative abundance of any absent species is 0. Consider the following notations: $a_{mn}$ is the number of shared species between the two areas *m* and *n*, $b_{mn}$ the number of species in *m* that are not present in *n* and $c_{mn}$ the number of species in *n* that are not present in *m*. With these notations, $S_m=a_{mn}+b_{mn}$. The relative abundance of any present species in area *m* is thus $1/(a_{mn}+b_{mn})$.

Similarly, the relative abundance of any present species in area $n$ is thus $1/(a_{mn}+c_{mn})$. If species are maximally dissimilar then eq. S4 reduces to

$$Q_{\beta,spe.div} = \sum_{m=1}^{M} w_m \sum_{n=1,n\neq m}^{M} w_n \left( \frac{1}{2}\sum_{i=1}^{S_\gamma} p_{im}^2 + \frac{1}{2}\sum_{i=1}^{S_\gamma} p_{in}^2 - \sum_{i=1}^{S_\gamma} p_{im} p_{in} \right) \quad \text{(S5)}$$

which means that the dissimilarity between areas $m$ and $n$ is considered to be

$$\left( \frac{1}{2}\sum_{i=1}^{S_\gamma} p_{im}^2 + \frac{1}{2}\sum_{i=1}^{S_\gamma} p_{in}^2 - \sum_{i=1}^{S_\gamma} p_{im} p_{in} \right) \quad \text{(S6)}$$

The following equalities hold, by definition of $a_{mn}$, $b_{mn}$ and $c_{mn}$, if all species present in an area also have identical abundances:

$$\sum_{i=1}^{S_\gamma} p_{im}^2 = (a_{mn}+b_{mn})\frac{1}{(a_{mn}+b_{mn})^2} = \frac{1}{(a_{mn}+b_{mn})} \quad \text{(S7)}$$

$$\sum_{i=1}^{S_\gamma} p_{in}^2 = (a_{mn}+c_{mn})\frac{1}{(a_{mn}+c_{mn})^2} = \frac{1}{(a_{mn}+c_{mn})} \quad \text{(S8)}$$

$$\sum_{i=1}^{S_\gamma} p_{im} p_{in} = a_{mn}\frac{1}{(a_{mn}+b_{mn})}\frac{1}{(a_{mn}+c_{mn})} \quad \text{(S9)}$$

It follows that if all species present in an area have identical abundances

$$\left( \frac{1}{2}\sum_{i=1}^{S_\gamma} p_{im}^2 + \frac{1}{2}\sum_{i=1}^{S_\gamma} p_{in}^2 - \sum_{i=1}^{S_\gamma} p_{im} p_{in} \right) = \frac{\frac{1}{2}(a_{mn}+c_{mn}) + \frac{1}{2}(a_{mn}+b_{mn}) - a_{mn}}{(a_{mn}+b_{mn})(a_{mn}+c_{mn})} = \frac{\frac{1}{2}(b_{mn}+c_{mn})}{(a_{mn}+b_{mn})(a_{mn}+c_{mn})}$$
(S10)

and thus

$$Q_{\beta,spe.ric} = \sum_{m=1}^{M} w_m \sum_{n=1,n\neq m}^{M} w_n \left( \frac{\frac{1}{2}b_{mn} + \frac{1}{2}c_{mn}}{\left(a_{mn}+b_{mn}\right)\left(a_{mn}+c_{mn}\right)} \right) \quad \text{(S11)}$$

The formula of $Q_{\beta,spe.ric}$ thus contains an index of the dissimilarity between areas $m$ and $n$ that we name $Q_{mn}$:

$$Q_{mn} = \left( \frac{\frac{1}{2}b_{mn} + \frac{1}{2}c_{mn}}{\left(a_{mn}+b_{mn}\right)\left(a_{mn}+c_{mn}\right)} \right) \quad \text{(S12)}$$

The dissimilarity $Q_{mn}$ between the two areas $m$ and $n$ reaches its maximum (1) when $a_{mn}$=0 and $b_{mn}$=$c_{mn}$=1, that is when each area contains only a unique, distinct species. This dissimilarity $Q_{mn}$ rapidly decreases when $a_{mn}$ departs (increases) from 0 (i.e., if species are shared between the sites) and/or when $b_{mn}$ and $c_{mn}$ depart (increase) from 1 (i.e., when each area has more than one species, be they shared or not with other areas). $Q_{\beta,spe.ric}$ can thus be viewed as the proximity to a case of complete species clustering where there are as many areas as species, each area containing a single species and areas do not share species.

#### S3. About $D_{\beta,spe.ric}$

To develop a component of community-to-community dissimilarity derived from $D_\beta$, we need to consider only two evenly weighted communities named *m* and *n*. In addition, a dissimilarity coefficient has to be zero between an entity (here a community) and itself ("hollow property"). Because the minimum of $D_\beta$ is 1 rather than 0, we obtained the following dissimilarity coefficient between plots *m* and *n* by applying $D_\beta$-1 to two evenly weighted communities *m* and *n*, considering maximally-dissimilar species that have even abundance within each community:

$$D_{mn} = \frac{b_{mn} + c_{mn}}{4a_{mn} + b_{mn} + c_{mn}} \quad \text{(S13)}$$

Indeed, if species are maximally dissimilar then $D_\beta$ reduces to (Eq. 8 in the main text)

$$D_{\beta,spe.div} = \frac{\left(\sum_{m=1}^{M} w_m \sum_{i=1}^{S_\gamma} p_{im}^2\right)}{\sum_{i=1}^{S_\gamma}\left(\sum_{m=1}^{M} w_m p_{im}\right)^2} \quad \text{(S14)}$$

If only two areas (*m* and *n*) are considered and they are evenly weighted Eq. S14 reduces to:

$$D_{\beta,spe.div} = \frac{\left(\frac{1}{2}\sum_{i=1}^{S_\gamma} p_{im}^2 + \frac{1}{2}\sum_{i=1}^{S_\gamma} p_{in}^2\right)}{\sum_{i=1}^{S_\gamma}\left(\frac{1}{2} p_{im} + \frac{1}{2} p_{in}\right)^2} \quad \text{(S15)}$$

If, in addition, species have even abundances in each area, then considering the same notations as in section S2 yields

$$D_{\beta,spe.ric} = \frac{\left(\tfrac{1}{2}\frac{1}{S_m} + \tfrac{1}{2}\frac{1}{S_n}\right)}{a_{mn} \times \left(\tfrac{1}{2}\frac{1}{S_m} + \tfrac{1}{2}\frac{1}{S_n}\right)^2 + b_{mn} \times \left(\tfrac{1}{2}\frac{1}{S_m}\right)^2 + c_{mn} \times \left(\tfrac{1}{2}\frac{1}{S_n}\right)^2} \quad \text{(S16)}$$

A species dissimilarity index, relying on species presence/absence data, can thus be defined as

$$D_{mn} = D_{\beta,spe.ric} - 1 = \frac{2\left(\frac{1}{a_{mn}+b_{mn}} + \frac{1}{a_{mn}+c_{mn}}\right)}{a_{mn}\times\left(\frac{1}{a_{mn}+b_{mn}} + \frac{1}{a_{mn}+c_{mn}}\right)^2 + b_{mn}\times\left(\frac{1}{a_{mn}+b_{mn}}\right)^2 + c_{mn}\times\left(\frac{1}{a_{mn}+c_{mn}}\right)^2} - 1$$

(S17)

$$D_{mn} = \frac{2\left(\frac{2a_{mn}+b_{mn}+c_{mn}}{(a_{mn}+b_{mn})(a_{mn}+c_{mn})}\right)}{a_{mn}\times\left(\frac{2a_{mn}+b_{mn}+c_{mn}}{(a_{mn}+b_{mn})(a_{mn}+c_{mn})}\right)^2 + b_{mn}\times\left(\frac{a_{mn}+c_{mn}}{(a_{mn}+b_{mn})(a_{mn}+c_{mn})}\right)^2 + c_{mn}\times\left(\frac{a_{mn}+b_{mn}}{(a_{mn}+b_{mn})(a_{mn}+c_{mn})}\right)^2} - 1$$

(S18)

$$D_{mn} = \frac{2(2a_{mn}+b_{mn}+c_{mn})(a_{mn}+b_{mn})(a_{mn}+c_{mn})}{a_{mn}\times(a_{mn}+b_{mn}+a_{mn}+c_{mn})^2 + b_{mn}\times(a_{mn}+c_{mn})^2 + c_{mn}\times(a_{mn}+b_{mn})^2} - 1 \quad \text{(S19)}$$

$$D_{mn} = \frac{2(2a_{mn}+b_{mn}+c_{mn})(a_{mn}+b_{mn})(a_{mn}+c_{mn})}{a_{mn}(a_{mn}+b_{mn})^2 + 2a_{mn}(a_{mn}+b_{mn})(a_{mn}+c_{mn}) + a_{mn}(a_{mn}+c_{mn})^2 + b_{mn}(a_{mn}+c_{mn})^2 + c_{mn}(a_{mn}+b_{mn})^2} - 1$$

(S20)

$$D_{mn} = \frac{2(2a_{mn}+b_{mn}+c_{mn})(a_{mn}+b_{mn})(a_{mn}+c_{mn})}{(a_{mn}+c_{mn})(a_{mn}+b_{mn})^2 + 2a_{mn}(a_{mn}+b_{mn})(a_{mn}+c_{mn}) + (a_{mn}+b_{mn})(a_{mn}+c_{mn})^2} - 1 \quad \text{(S21)}$$

$$D_{mn} = \frac{2(2a_{mn}+b_{mn}+c_{mn})}{(a_{mn}+b_{mn}) + 2a_{mn} + (a_{mn}+c_{mn})} - 1 = \frac{2(2a_{mn}+b_{mn}+c_{mn})}{4a_{mn}+b_{mn}+c_{mn}} - 1 \quad \text{(S22)}$$

$$D_{mn} = \frac{4a_{mn}+2b_{mn}+2c_{mn}-4a_{mn}-b_{mn}-c_{mn}}{4a_{mn}+b_{mn}+c_{mn}} = \frac{b_{mn}+c_{mn}}{4a_{mn}+b_{mn}+c_{mn}} \quad \text{(S23)}$$

□

$D_{mn}$ varies between 0 and 1. If $a_{mn} = 0$ then $D_{mn} = 1$ whatever the values of $b$ and $c$. This means that if two plots do not share species they will be considered as maximally dissimilar as soon as they have one species each, or equivalently that two sites will be considered maximally dissimilar independently of the number of unshared species: two plots with no shared species and 1 species each will be considered as dissimilar as two plots with no shared species and say 1000 species each. If $b_{mn} = c_{mn} = 0$ then $D_{mn} = 0$ whatever the number of shared species ($a_{mn}$).

The index $D_{mn}$ belongs to the same family of dissimilarity indices as Sørensen dissimilarity index

$$\text{Sørensen}_{mn} = \frac{b_{mn} + c_{mn}}{2a_{mn} + b_{mn} + c_{mn}} \qquad \text{(S24)}$$

Jaccard dissimilarity index

$$Jaccard_{mn} = \frac{b_{mn} + c_{mn}}{a_{mn} + b_{mn} + c_{mn}} \qquad \text{(S25)}$$

and Sokal & Sneath (1963) dissimilarity index (see also Anderberg, 1973; and S5 coefficient of Gower & Legendre, 1986)

$$SokalSneath_{mn} = \frac{b_{mn} + c_{mn}}{\frac{1}{2}a_{mn} + b_{mn} + c_{mn}} \qquad \text{(S26)}$$

### S4. The values of $U_R$ tend to unity under the random scenario as the number of individuals per area increases (Appendix S1)

As we observed that UR was dependent on the number of individuals per community in our simulations, we also considered communities composed of 25000 individuals to analyze its values (Fig. S1).

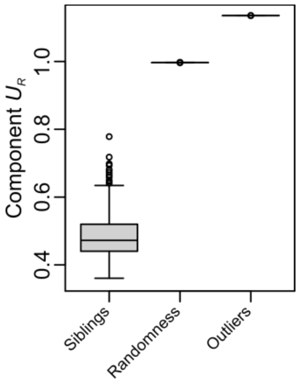


**FIGURE S1** Distribution of component $U_R$ (ratio of $D_\beta$ to $D_{\beta,spe.div}$) over simulations summarized by box plots considering 25000 individuals in each area. Box plots are given for three scenarios: Siblings (Scenario 5 in section *Case Studies*), Randomness (Scenario 1 in section *Case Studies*) and Outliers (Scenario 4 in section *Case Studies*).

**Supplementary References**

Anderberg, M. R. 1973. Cluster Analysis for Applications. New York: Academic Press.

Gower, J. C., and P. Legendre. 1986. Metric and Euclidean properties of dissimilarity coefficients. *Journal of Classification* 3: 5–48. https://doi.org/10.1007/bf01896809

Rao, C. R. 1982. Diversity and dissimilarity coefficients: a unified approach. *Theoretical Population Biolog*y 21: 24–43. https://doi.org/10.1016/0040-5809(82)90004-1

Sokal, R. R., and P. H. A. Sneath. 1963. Principles of Numerical Taxonomy. San Francisco: Freeman.
